# VEGABrain: a Visualizer of Enriched Gene Analysis for the developing human brain

**DOI:** 10.64898/2026.11.19.767876

**Authors:** Isaac Medeiros, João V. F. Cavalcante, Diego Marques-Coelho

**Affiliations:** Bioinformatics Multidisciplinary Environment, Federal University of Rio Grande do Norte, Rua do Horto, Natal, 59076-550, Rio Grande do Norte, Brazil

**Keywords:** Neuroinformatics, Brain development, BrainSpan, Gene set enrichment, ssGSEA, Data visualization, Web application

## Abstract

RNA sequencing datasets are widely reused resources in developmental neuroscience, yet inspecting them spatially remains unclear and difficult. Here, we present VEGABrain, an openly accessible web tool that renders bulk RNA-seq onto brain cortical and subcortical atlas surfaces across five human developmental windows spanning the first trimester to adulthood. Beyond single-gene maps, VEGABrain contributes a composite per-region enrichment layer using single-sample gene set enrichment analysis (ssGSEA) scores precomputed for 18,650 gene sets drawn from Gene Ontology, Reactome, KEGG, BioCarta, the Human Phenotype Ontology and the MSigDB hallmark collection, and aggregated to a score for every gene set, developmental window and brain region. Furthermore, VEGABrain allows users to submit a custom gene list and obtain its ssGSEA map across development, computed on demand. Submitted lists can also be tested: two complementary per-area tests—one competitive on the ssGSEA score, one for over-representation of area-specific marker genes—mark the regions where a list stands out, and a developmental-stage test reports the windows in which it peaks, for the whole brain and for six major regions. VEGABrain is open source and requires no installation, offering easy-to-query expression throughout development and brain areas.

## 1 Introduction

The transcriptome of the developing human brain is one of the most heavily reused datasets in neuroscience. The BrainSpan atlas (BrainSpan 2011; Kang et al. 2011; Miller et al. 2014) sampled bulk tissue from cortical and subcortical structures across prenatal and postnatal donors, and subsequent integrative work built on it to relate developmental expression to neuropsychiatric risk (Li et al. 2018) and to evolutionary divergence between human and macaque (Zhu et al. 2018).

What has not kept pace is the ease of seeing it. A researcher who wants to know where and when a gene, or a set of genes, is expressed in the developing brain has essentially two options today, and both impose costs that are unrelated to the biological question.

The first is the resource portal. Portals in this family—BrainSpan itself, and the wider family of Allen Institute browsers (Lau et al. 2008; Sunkin et al. 2013)—return the data as tables, line plots or heat maps indexed by structure acronym. This, unfortunately, is not anatomical: the reader must hold the mapping from DFC, MFC or STC to a location in the brain, and a heat map of 26 structure codes does not convey a spatial gradient the way a rendered cortex does. It is also gene-at-a-time. There is no route from a gene set, a pathway, or the output of a differential-expression analysis to a spatial view. Newer hosted portals have narrowed part of this gap: BrainScope links genes and samples through t-SNE maps to choropleths over anatomical slices (Huisman et al. 2017), and SMDB browses spatial transcriptomic data in two and three dimensions (Cao et al. 2023). Neither, however, scores a gene set across a human developmental parcellation (Section 4.1).

The second option would be through brain-plotting tools as—ggseg and ggseg3d (Mowinckel and Vidal-Piñeiro 2020) and cerebroViz (Bahl et al. 2017) in R, neuromaps in Python (Markello et al. 2022), and abagen for the adult Allen Human Brain Atlas (Hawrylycz et al. 2012; Markello et al. 2021). But that route presupposes a user who writes R or Python, who is willing to download and hold a 52,376*×*524 expression matrix locally, and who will construct and validate the mapping from BrainSpan’s tissue nomenclature onto an atlas parcellation. Each of those steps is a place where a project stalls or a mapping silently diverges between laboratories (Markello et al. 2021).

Additionally, modern analyses rarely end with one gene. They end with a list, be it a differentially expressed set, a module from a co-expression network or a panel of risk genes for a disorder. The natural question is then not “where is gene *X* expressed?” but “where and when is this programme active?”. Answering it requires collapsing many genes into a single per-sample summary. Single-sample gene set enrichment analysis (ssGSEA) (Barbie et al. 2009), a per-sample variant of GSEA (Subramanian et al. 2005) implemented alongside GSVA (Hänzelmann et al. 2013), provides exactly such a composite score, and has been used as a quantitative readout of pathway activity in expression data (Yi et al. 2020). The incorporation of composite multigene activity scores (such as those generated by ssGSEA or GSVA) into anatomical models directly addresses the challenge of condensing high-dimensional expression profiles into interpretable summaries, ensuring noise reduction and dimensionality reduction compared to classical single-gene-based analyses (Hänzelmann et al. 2013). To our knowledge no tool has applied it as a *spatial* layer over the developing human brain.

VEGABrain (Visualizer of Enriched Gene Analysis) addresses these three gaps in one interface, reachable at https://mcblab.github.io/VEGABrain/ with no installation, account or download. It maps BrainSpan samples onto cortical and subcortical atlas parcels across five developmental windows; it adds a composite enrichment layer of ssGSEA scores precomputed for 18,650 gene sets from the Gene Ontology (Ashburner et al. 2000; The Gene Ontology Consortium et al. 2023), Reactome (Gillespie et al. 2022), KEGG (Kanehisa and Goto 2000), BioCarta, the Human Phenotype Ontology (Köhler et al. 2021) and the MSigDB hallmark collection (Liberzon et al. 2011, 2015); and it scores user-submitted gene lists on demand, closing the loop from an enrichment result (Chen et al. 2013; Kuleshov et al. 2016) back to anatomy. A submitted list may also carry log2 fold-change values from an external comparison—a different disease model, treatment or condition—which are then mapped against the BrainSpan expression data, so that analyses can follow directions the dataset was not originally built for, not just the comparisons already included in it. A submitted list is also tested statistically: VEGABrain marks the areas where it stands out against random genes of the same expression level, and reports the developmental windows in which it peaks, so that a map can be read as a result rather than only as a picture. Both the maps and the numbers behind them are exportable, so that VEGABrain is a starting point for downstream analysis, consistent with reusable research data (Wilkinson et al. 2016).

## 2 Materials and Methods

### 2.1 Expression data and preprocessing

VEGABrain is built on the BrainSpan Developmental Transcriptome RNA-Seq release, Gencode v10 RPKM values summarised to genes (BrainSpan 2011; Li et al. 2018). The distributed matrix contains 52,376 gene rows and 524 sample columns. Rows with duplicate HGNC symbols are dropped, keeping the first occurrence, and values are transformed as log_2_(RPKM + 1). No further normalisation or batch correction is applied; the released RPKM values are used as the provider distributes them, so that scores computed here remain comparable with analyses performed directly on the public matrix. Preprocessing and the scoring described in Section 2.4 are performed once, offline (Figure 1).

**Fig. 1.**
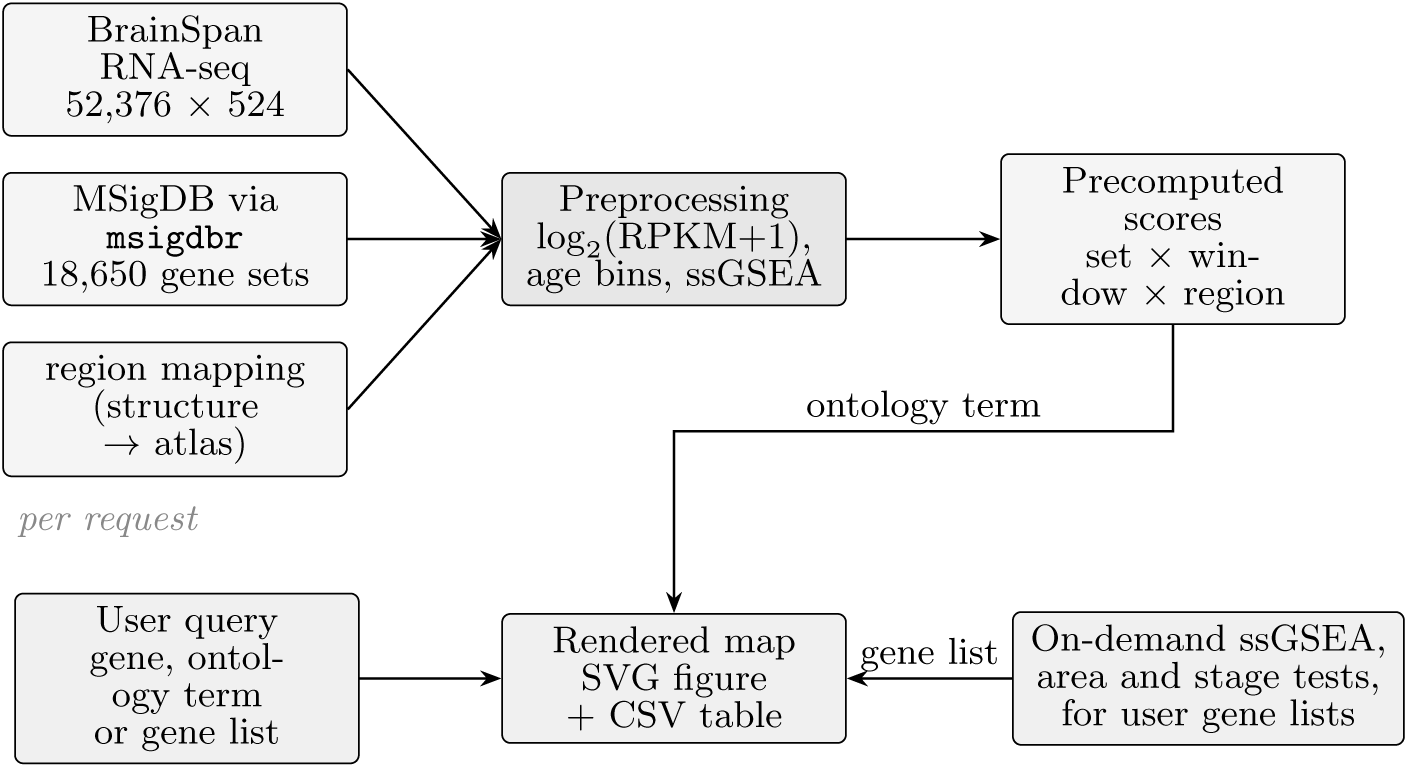
VEGABrain data flow. The top row runs once, offline; the bottom row runs per request. Ontology queries are served from the precomputed score table, whereas a user-submitted gene list triggers a fresh ssGSEA run over all 524 samples and, on request, the area and stage tests of Section 2.5 against 4,000 expression-matched random gene sets.

### 2.2 Developmental windows

BrainSpan donors span 31 distinct ages from 8 post-conceptional weeks (pcw) to 40 years. Individual ages are too sparsely sampled to render as separate maps, so donors are binned into five developmental windows: first trimester (8–12 pcw, *n* = 5 donors), second trimester (13–21 pcw, *n* = 10), third trimester (24–37 pcw, *n* = 5), child (*n* = 14) and adult (*n* = 8).

The two postnatal windows split the postnatal donors at 18 years, where BrainSpan’s sampling has a natural gap: no donor is aged between 15 and 18 years. The child window therefore holds every postnatal donor under 18: 14 donors aged 4 months to 15 years, five of them under two years, four aged 2 to 4 years and five aged 8 to 15 years. It spans infancy, childhood and adolescence (periods 8–12 of Kang et al. 2011), so a result in this window describes the maturing postnatal brain as a whole rather than any one of those stages. The adult window holds the eight donors aged 18 to 40 years, one per age, four male and four female. No donor is older than 40, so this window covers young adulthood and says nothing about ageing.

The donor counts and age ranges are also given in Table 1; the donor count is displayed in every facet label in the interface so that the reader is never shown a map without also being shown how much evidence stands behind it.

**Table 1.** Data underlying VEGABrain.

| Property | Value |
|---|---|
| Expression source | BrainSpan RNA-Seq, Gencode v10 RPKM, gene-summarised |
| Matrix dimensions | 52,376 genes × 524 samples |
| Transformation | log<sub>2</sub>(RPKM + 1); duplicate symbols dropped |
| Developmental windows | 5 (see below) |
| 1st trimester | 5 donors, 8–12 pcw |
| 2nd trimester | 10 donors, 13–21 pcw |
| 3rd trimester | 5 donors, 24–37 pcw |
| Child | 14 donors, 4 months–15 years |
| Adult | 8 donors, 18–40 years |
| Atlas parcels (micro) | 28 (21 Desikan–Killiany cortical + 7 aseg) |
| Anatomical groups (macro) | 6 (frontal, parietal, temporal, occipital, subcortical, cerebellum) |
| Views per map | 4 (DK lateral, DK medial, aseg sagittal, aseg coronal) |
| Gene sets scored | 18,650 total |
| GO biological process | 7,538 |
| Human Phenotype Ontology | 5,793 |
| GO molecular function | 1,872 |
| Reactome | 1,839 |
| GO cellular component | 1,080 |
| BioCarta | 292 |
| KEGG (legacy) | 186 |
| MSigDB hallmark | 50 |

### 2.3 Mapping BrainSpan structures to an atlas

BrainSpan labels tissue by dissected structure (for example *dorsolateral prefrontal cortex* ), whereas ggseg draws atlas parcels (for example *rostral middle frontal* ). Cortical regions map onto the Desikan–Killiany parcellation (Desikan et al. 2006) and subcortical structures onto the FreeSurfer aseg segmentation (Fischl et al. 2002), both accessed through ggseg (Mowinckel and Vidal-Piñeiro 2020).

Each atlas parcel takes the samples of one BrainSpan structure, except the thalamus and the cerebellum, which pool the two names BrainSpan gives each (dorsal thalamus and mediodorsal nucleus of the thalamus; cerebellum and cerebellar cortex). BrainSpan samples far fewer structures than the atlases draw, so the assignment is many-to-one: 22 of BrainSpan’s 26 structures cover 28 atlas parcels—21 of the 34 Desikan–Killiany cortical parcels and seven aseg structures (amygdala, hippocampus, thalamus, caudate, putamen, nucleus accumbens and cerebellum)—and parcels that fall within one dissected structure share its value (for example, the rostral and caudal middle frontal parcels both take the dorsolateral prefrontal cortex samples, and the caudate, putamen and nucleus accumbens all take the striatum samples). The four structures left out, the medial, lateral and caudal ganglionic eminences and the upper rhombic lip, are transient fetal structures with no counterpart in either atlas. Atlas parcels with no BrainSpan counterpart are left unassigned and drawn in grey: 13 Desikan–Killiany parcels, among them the superior frontal gyrus, insula, precuneus and fusiform gyrus, and aseg structures such as the brain stem, pallidum and choroid plexus. The complete assignment is a single table, provided as Supplementary File 1 and read by the application as is, so it can be inspected and changed rather than reimplemented per project.

The interface exposes two levels of granularity. At the *micro* scale each of the 28 mapped atlas parcels carries its own value. At the *macro* scale regions are pooled into six anatomical groupings—frontal, parietal, temporal and occipital lobes, subcortical structures and cerebellum—the mean is taken within each grouping, and that mean is painted back onto every region belonging to it. The macro scale exists because BrainSpan sampling density varies sharply across structures: pooling trades spatial resolution for donor support, and letting the user switch between the two makes that trade-off explicit rather than hidden. Regions with no corresponding sample are drawn in grey rather than imputed.

Every map is rendered as four stacked panels—right lateral cortex, right medial cortex, subcortical sagittal and subcortical coronal—each faceted into the five developmental windows, giving a single figure in which spatial and temporal structure are read on orthogonal axes.

### 2.4 Composite enrichment scores

For a gene set *G* and a sample *j*, ssGSEA (Barbie et al. 2009) ranks all genes by their expression within that sample and computes an enrichment score from the normalised difference between the empirical cumulative distribution of the genes in *G* and that of the genes outside it:

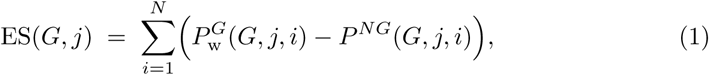

where *i* indexes genes in decreasing order of expression in sample *j*, *P_w_^G^* is the weighted cumulative distribution over genes in the set and *P ^NG^* the cumulative distribution over the rest. Because the score is computed from within-sample ranks, it is invariant to the monotone transformation applied in Section 2.1 and does not require a reference group or a phenotype contrast—which is what makes it usable here, where each sample is a single donor–structure pair rather than a member of a case/control design. We use the ssgseaParam interface of the GSVA package (Hänzelmann et al. 2013) with default parameters.

Gene sets are retrieved through msigdbr (Dolgalev 2026) for *Homo sapiens* and comprise the MSigDB hallmark collection (Liberzon et al. 2015), the canonical pathway subcollections BioCarta, KEGG (Kanehisa and Goto 2000) and Reactome (Gillespie et al. 2022), and the full C5 ontology collection, which supplies Gene Ontology biological process, molecular function and cellular component terms (Ashburner et al. 2000; The Gene Ontology Consortium et al. 2023) together with Human Phenotype Ontology terms (Köhler et al. 2021). The resulting 18,650 sets are broken down by collection in Table 1.

Scores are computed once for every gene set against all 524 samples, then aggregated: for each (gene set, developmental window, region) triple the mean score over the contributing samples is stored, and analogously for macro-regions, so that an ontology query needs no enrichment run at request time. When instead a user submits their own gene list, the symbols are validated against the matrix (unmatched symbols are reported back rather than silently dropped), ssGSEA is run for that single set across all samples, and the same aggregation is applied before rendering.

### 2.5 Significance testing for gene lists

A map shows where a score is high; it does not say whether that is more than chance. For a submitted list, VEGABrain therefore offers three tests, run on request and reported on the map and in the exported table (Table 2). All three are computed from the expression matrix and its sample annotation at request time, with no precomputed reference. The unit of inference is always the BrainSpan structure or region class, never the atlas parcel: results are painted onto parcels only for display, because parcels fed by the same structure carry the same measurement (Section 2.3).

**Table 2.**
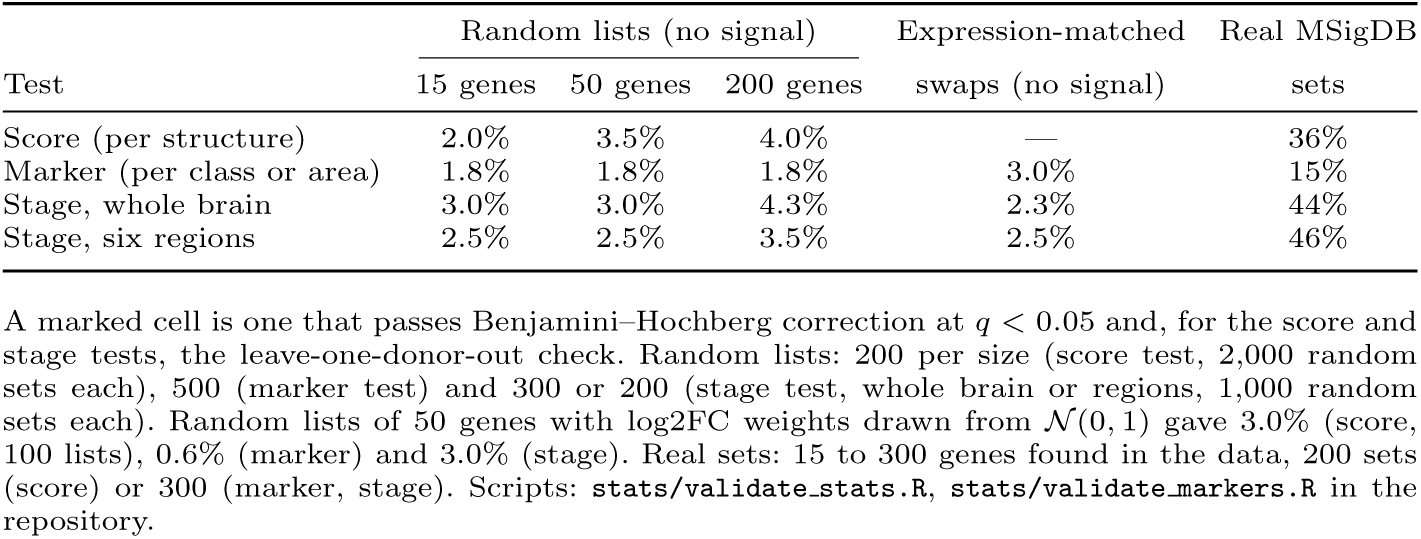
Calibration of the significance tests: share of lists with at least one marked cell.

|  | Random lists (no signal) |  |  | Expression-matched | Real MSigDB |
|---|---|---|---|---|---|
| Test | 15 genes | 50 genes | 200 genes | swaps (no signal) | sets |
| Score (per structure) | 2.0% | 3.5% | 4.0% | — | 36% |
| Marker (per class or area) | 1.8% | 1.8% | 1.8% | 3.0% | 15% |
| Stage, whole brain | 3.0% | 3.0% | 4.3% | 2.3% | 44% |
| Stage, six regions | 2.5% | 2.5% | 3.5% | 2.5% | 46% |

A marked cell is one that passes Benjamini–Hochberg correction at *q* < 0.05 and, for the score and stage tests, the leave-one-donor-out check. Random lists: 200 per size (score test, 2,000 random sets each), 500 (marker test) and 300 or 200 (stage test, whole brain or regions, 1,000 random sets each). Random lists of 50 genes with log2FC weights drawn from $\mathcal{N}(0, 1)$ gave 3.0% (score, 100 lists), 0.6% (marker) and 3.0% (stage). Real sets: 15 to 300 genes found in the data, 200 sets (score) or 300 (marker, stage). Scripts: stats/validate\_stats.R, stats/validate\_markers.R in the repository.

#### Score test

It asks whether the list, taken as a whole, is more active in a structure than in the other structures sampled in the same window. For every structure–window cell with at least three donors (70 of 87 cells), the list’s mean ssGSEA score is taken as its deviation from the mean over that window’s structures, and this deviation is ranked against those of 4,000 random gene sets of the same size, each gene replaced by one from the same bin of mean expression (50 bins over the MSigDB gene universe). Centring within the window is what makes the test spatial: without it, any list with a developmental trend would mark every structure of its peak window. *p*-values are corrected by the Benjamini–Hochberg procedure over all tested cells (Benjamini and Hochberg 1995), and a cell is reported only if it stays below 0.05 when each of its donors is left out in turn. For a list with log2FC values the score is the weight-averaged *z*-score of its genes, and the random sets carry the same weights. So that 4,000 random sets can be scored per request, ssGSEA is computed from per-sample rank tables built once; this reproduces GSVA to a relative error below 2 *×* 10*^−^*^5^.

#### Marker test

It asks whether the list contains the genes that set an area apart, following the logic of tissue- and cell-type-specific expression analysis (Dougherty et al. 2010; Xu et al. 2014). In each window, limma (Ritchie et al. 2015) fits expression on structure with donor as a fixed effect, so that every contrast is made within donors, and tests two levels of contrast: each of six region classes (neocortex, hippocampus, amygdala, striatum, thalamus, cerebellum) against the equally weighted mean of the other five, and each neocortical area against the mean of the other neocortical areas. Genes with *q <* 0.05, log_2_ fold change *≥* 1 and mean log_2_(RPKM + 1) *≥* 1 are that area’s markers. A one-sided Fisher exact test then compares the list with the markers, within the window’s expressed MSigDB genes, with Benjamini–Hochberg correction over all tested cells. A cell is tested if it has at least 10 markers and three informative donors (36 cells), and a hit needs *q <* 0.05 and at least two shared genes. A list with log2FC values is tested separately for its up- and down-regulated genes. Unlike the score test, the marker test names the user’s genes that are responsible for each hit.

#### Stage test

It asks in which window, rather than in which area, a list stands out—a question the two area tests cannot answer, because both compare areas within one window. For each window, the list’s ssGSEA score is averaged by structure, then by region class, then over the six region classes of the PsychENCODE grouping (NCX, HIP, AMY, STR, MD, CBC; Li et al. 2018) with equal weight, so that every window has the same regional make-up even though BrainSpan sampled different structures at different ages. The window value minus the list’s mean over the five windows is ranked against the same 4,000 expression-matched random sets; the test is two-sided, Benjamini–Hochberg corrected over the windows, and guarded by leaving out each donor of the window in turn. It is repeated inside each region class, where a region–window cell is tested only if it has at least three donors; in the third trimester only the neocortex and the cerebellum qualify. The whole-brain and regional rows are corrected as separate families, and a list with log2FC values is also tested as its up- and down-regulated halves.

#### Calibration

Each test was run on lists that carry no signal (Table 2): uniform random lists of 15, 50 and 200 genes, random lists with log2FC weights, and real MSigDB sets with every gene swapped for one of matched expression. At most 5% of such lists produced a single marked cell in any test. On real MSigDB sets, by contrast, 36% had a score-test hit, 15% a marker-test hit and 44% a stage-test hit—the last reflecting that development is the dominant axis of variation in brain expression, which is why the stage test reports an effect size (*z*) for every window. Positive controls behaved as expected: eight striatal markers (*PPP1R1B*, *DRD1*, *DRD2* and others) mapped to the striatum in all four testable windows and nowhere else, and the astrocyte-development, myelin-assembly and cell-cycle sets (GOBP ASTROCYTE DEVELOPMENT, GOBP MYELIN ASSEMBLY, HALLMARK G2M CHECKPOINT and HALLMARK E2F TARGETS) peaked in the third trimester, after birth and in the first two trimesters, respectively. Every list used in this work is given in Supplementary Table 2.

### 2.6 Implementation

VEGABrain is an R (R Core Team 2026) service built with plumber (Schloerke and Allen 2026) behind a browser interface (Figure 1). Maps are rendered with ggplot2 (Wickham 2016) and ggseg (Mowinckel and Vidal-Piñeiro 2020), and can be downloaded as vector (SVG) figures together with the per-region values behind them as CSV. The same service is exposed as a public HTTP API, documented in the source repository, so that maps and tables can also be retrieved programmatically. The significance tests (Section 2.5) use limma (Ritchie et al. 2015) for the marker contrasts and are exposed through the same API: a significance parameter on the gene-list map and table, and two endpoints that return every tested cell, POST /area enrichment and POST /stage enrichment. The reference tables the tests need are built from the expression matrix on the first request after the service starts (about 20 s); a list then takes from one to about 5 s, depending on its size and on whether it carries log2FC values. In the interface, the chosen area test marks the map, the stage result is shown beside it, and the exported table carries all three tests.

## 3 Results

VEGABrain offers three query modes—a single gene, a precomputed ontology term, or a user-submitted gene list—on a single working page. All three produce the same four-panel, five-window figure at either region granularity (Section 2.3), and a reference map labelled by region name is available so that unfamiliar atlas parcels can be identified. In gene-list mode the tests of Section 2.5 add marks to the map and a developmental-stage panel beside it. We illustrate each mode with a case whose developmental behaviour is independently established, so that the maps can also be judged for biological face validity.

### 3.1 Single-gene maps

Gene mode renders the mean log_2_(RPKM + 1) of any gene in the matrix per region and developmental window, on a fixed colour scale bounded at 6, so that maps for different genes are directly comparable rather than each being auto-scaled to its own range. Oligodendrocyte and myelination markers (*SOX10*, *MBP*, *PLP1* ) should be low prenatally and rise sharply in the child and adult windows as myelination proceeds; Figure 2 shows *SOX10*.

**Fig. 2.**
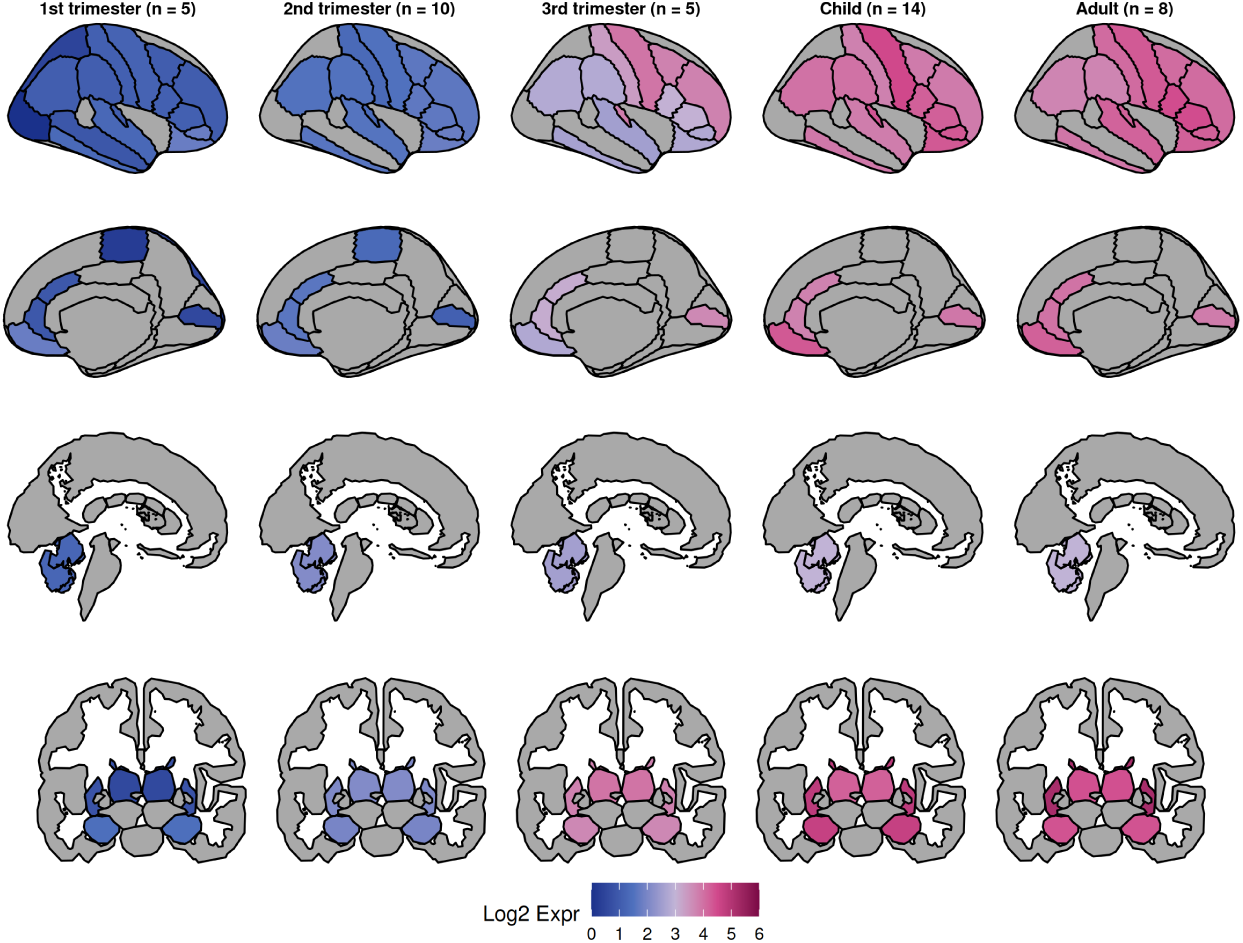
Single-gene mode. Mean log_2_(RPKM + 1) for *SOX10* across the five developmental windows (columns) in four views (rows): right lateral cortex and right medial cortex on the Desikan–Killiany parcellation, and subcortical structures in sagittal and coronal section on the aseg segmentation. Grey indicates regions with no BrainSpan sample in that window.

### 3.2 Gene-set maps

Ontology mode returns the mean ssGSEA score of any of the 18,650 precomputed gene sets per region and window (Figure 3). This is the composite layer described in Section 2.4, and it is what distinguishes VEGABrain from a gene-at-a-time browser: a single map summarises the coordinated behaviour of tens to hundreds of genes. The member genes of the selected set are listed alongside the map, so the composition of any score is inspectable. Astrocytes are generated after neurons, once cortical neurogenesis gives way to gliogenesis (Rowitch and Kriegstein 2010), so GOBP ASTROCYTE DEVELOPMENT should rise late in prenatal development. Its map peaks in the third trimester, in cortex and subcortex alike (Figure 3a). The composite score is informative here: the individual genes of the term are noisy across 28 sparsely sampled parcels, whereas the ssGSEA summary of its 42 genes is not. Submitting those genes in gene-list mode (Supplementary Table 2) tests the reading directly (Figure 3b): against random genes of the same expression level, the term is higher in the third trimester (whole-brain *z* = 6.1; neocortex 4.7, cerebellum 4.3) and lower in the first (*z* = *−*2.6). The hippocampus, amygdala, striatum and thalamus show the same third-trimester peak (*z* = 4.6 to 5.9) but cannot be tested there, as each was sampled from only two donors in that window. The example also shows why the stage test is needed. Because the term peaks everywhere at once, neither area test, both of which compare areas within a window, marks any area in the third trimester; the peak is a matter of when, not where.

**Fig. 3.**
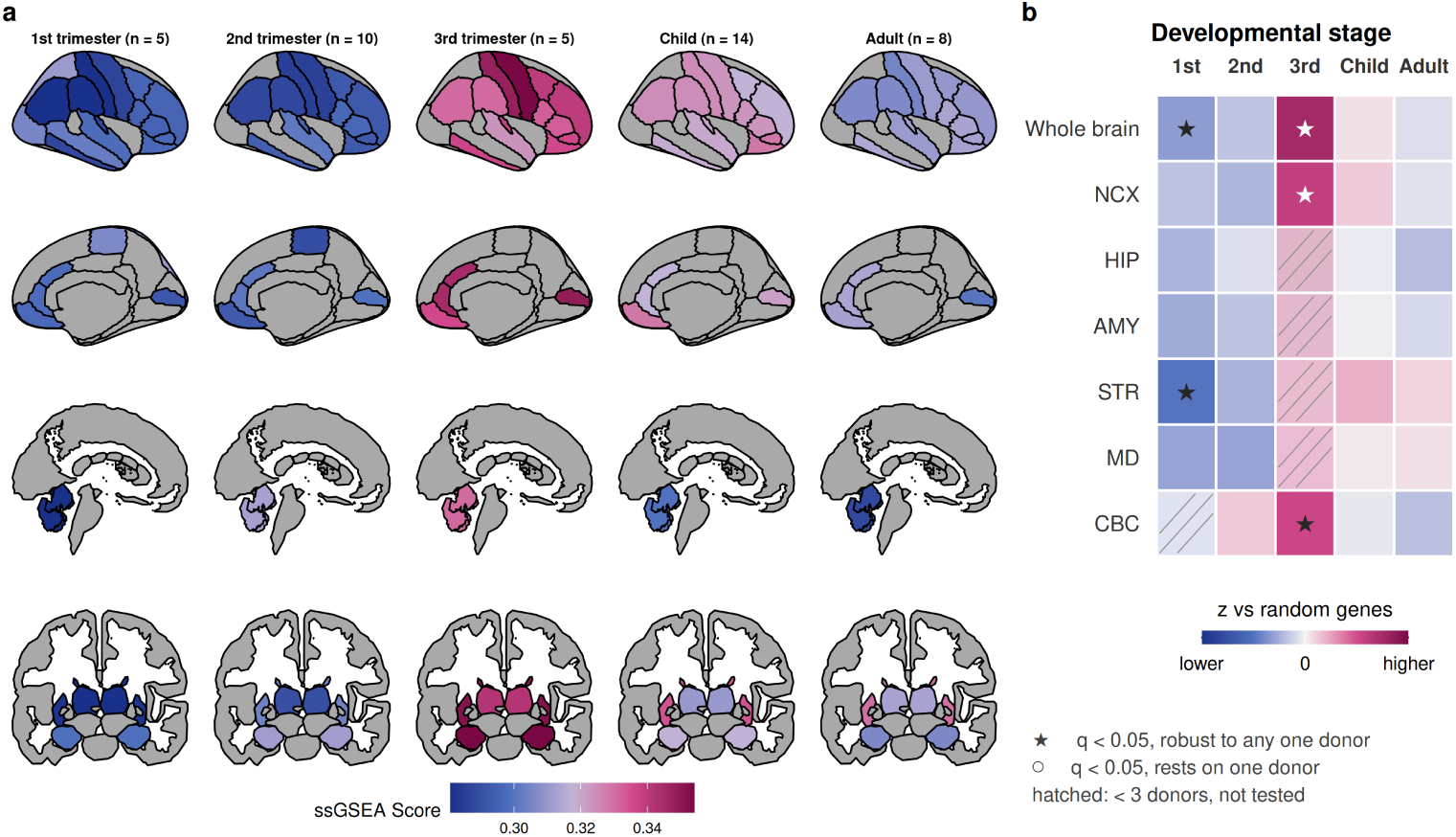
Ontology mode. (a) Composite ssGSEA score for GOBP ASTROCYTE DEVELOPMENT across development. The score summarises every gene in the term into one value per region and window; the same term can be viewed at micro granularity (28 atlas parcels) or macro granularity (six anatomical groupings). (b) Developmental-stage test of the term’s 42 genes (Section 2.5), for the whole brain and the six PsychENCODE regions. Colour is *z* against 4,000 expression-matched random gene sets, centred over the five windows; a star marks *q <* 0.05 that survives leaving out any one donor, a ring *q <* 0.05 resting on one donor, and hatching a cell with fewer than three donors, which is not tested. The term peaks in the third trimester in every region, but only the neocortex and the cerebellum have enough donors in that window to be tested.

### 3.3 User-submitted gene lists

In gene-list mode, a pasted list of symbols is validated and scored on demand (Figures 4 and 5). Symbols absent from the matrix are listed back to the user explicitly, so a map is never quietly computed on a subset of what was submitted. This is the path from an external result to anatomy: the output of a differential expression analysis, a co-expression module, or a disorder risk panel can be mapped across development directly, without the user preparing data or writing code. GABAergic interneuron markers (*GAD1*, *GAD2*, *SLC32A1* ) should be present from the second trimester onward with subcortical and then cortical distribution. In addition to entering genes, for each entry, the researcher can add a logFC value of their choice. The process can simulate how that variation in expression might impact the developmental process; the tests then treat the up- and down-regulated genes separately, since the two halves of a signature can belong to different areas or stages.

**Fig. 4.**
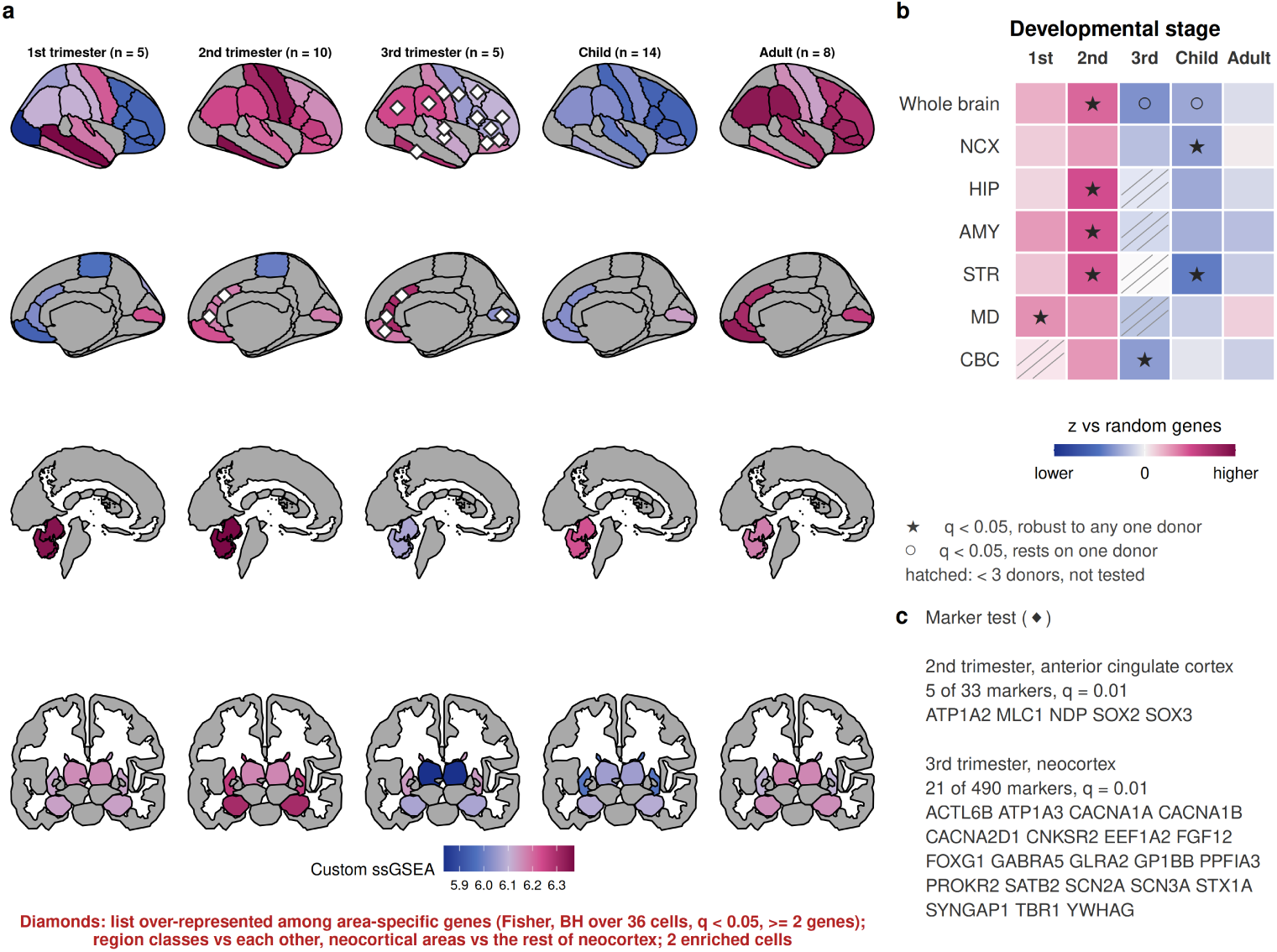
Where and when a gene list is enriched: the 262 genes of HP AUTISM found in BrainSpan. (a) ssGSEA map with the marker test: diamonds mark the areas whose specific genes are over-represented in the list (one-sided Fisher test against limma area markers, *q <* 0.05, at least two shared genes). A hit for the neocortex as a whole, here in the third trimester, marks every neocortical area sampled in that window. (b) Developmental-stage test, as in Figure 3b: the list peaks in the second trimester, as autism risk genes are known to (Willsey et al. 2013; Parikshak et al. 2013). (c) The cells marked in (a), with the shared genes.

**Fig. 5.**
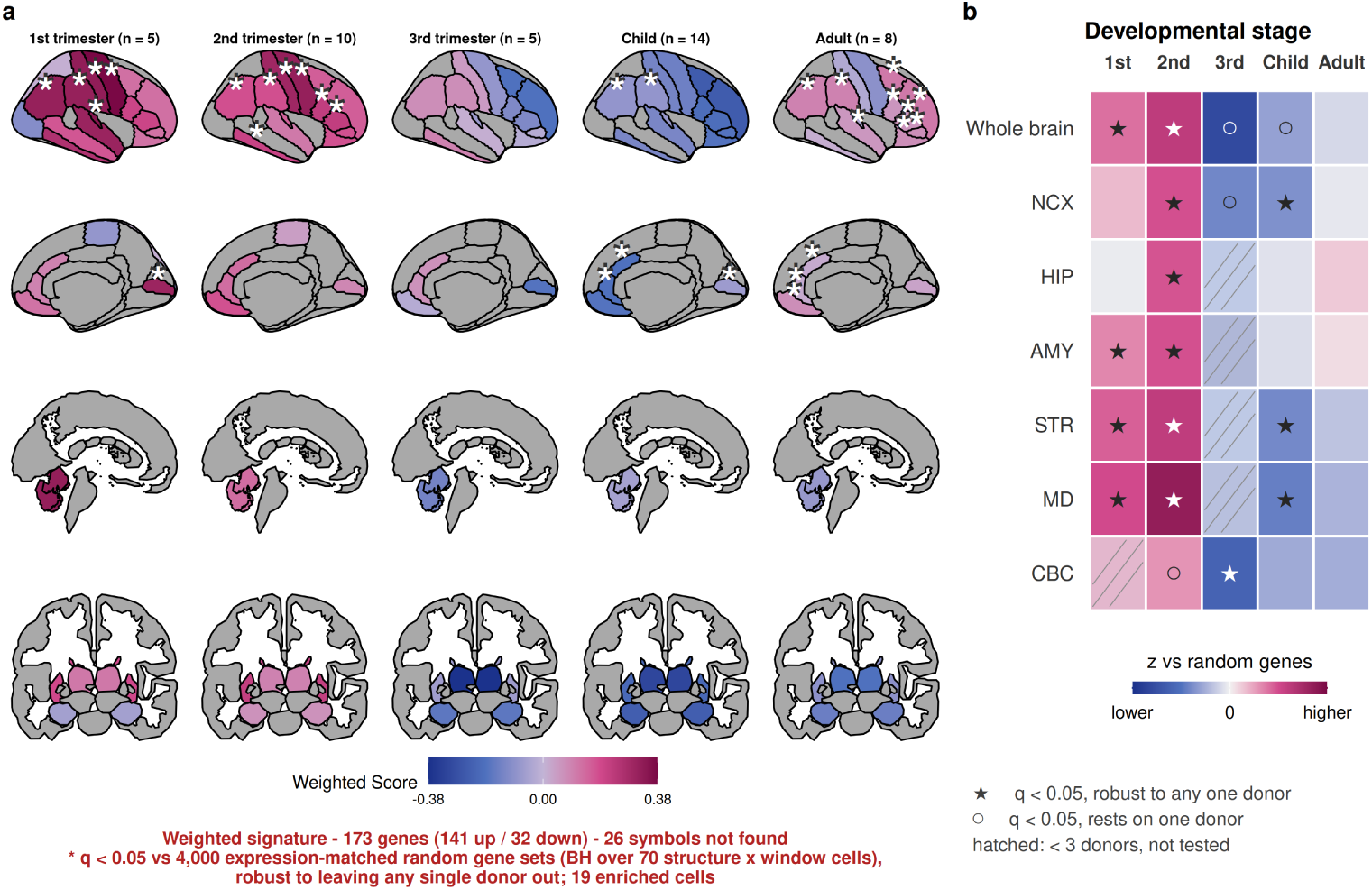
A signature with fold changes: the genes changed by 6 h of 50 mM ethanol in two-month-old human cerebral organoids (Arzua et al. 2020), submitted with their signed log2 fold changes (173 of 199 genes found in BrainSpan; 141 up, 32 down). The list is a published differential-expression result, mapped without local installation or coding. (a) Weighted score: magenta where the genes ethanol raised are high relative to those it lowered, blue where the reverse holds. Asterisks mark the score test: structures where the weighted score exceeds that of 4,000 expression-matched random gene sets carrying the same weights (*q <* 0.05, robust to leaving out any donor); all 19 are neocortical areas. (b) Developmental-stage test of the weighted list, as in Figure 3b: higher in the first and second trimesters, the stage that organoids of this age resemble.

### 3.4 Where and when: testing a gene list

Figure 4 applies the three tests to a list whose developmental answer is known from independent work: the 262 genes of the Human Phenotype Ontology term HP AUTISM that are present in BrainSpan (Supplementary Table 2). Autism risk genes converge on mid-fetal cortical development (Willsey et al. 2013; Parikshak et al. 2013). The marker test, drawn on the map (Figure 4a), asks which areas’ specific genes the list contains, and names them (Figure 4c). The list holds 21 of the 490 neocortex markers of the third trimester, among them *SATB2*, *FOXG1*, *TBR1* and *SYNGAP1*, so every neocortical area sampled in that window is marked; it also holds 5 of the 33 markers of the second-trimester anterior cingulate cortex, mostly progenitor and glial genes (*SOX2*, *SOX3*, *ATP1A2* ). The stage test recovers the timing from the list alone: the list peaks in the second trimester (whole-brain *z* = 3.4; hippocampus, amygdala and striatum, *z* = 3.6 to 3.9) and is lower in the third trimester and in the child window (Figure 4b). The score test, the alternative mark for the same map, finds ten neocortical cells, in the second trimester (primary motor and somatosensory cortex, inferolateral temporal cortex), in the child window and in adulthood. The term is broader than a curated risk list—it includes syndromic genes such as the Bardet–Biedl ciliopathy family—so the effect sizes are moderate, yet the three tests agree on a prenatal, cortical reading.

### 3.5 A signature with fold changes: ethanol in cerebral organoids

A list can also carry the log2 fold changes of a differential-expression result. Arzua et al. (2020) exposed two-month-old cerebral organoids, derived from human induced pluripotent stem cells, to 50 mM ethanol for 6 h, and report 199 genes changed more than twofold (*P <* 0.05), 162 up and 37 down. Submitting that table as published, with signed log2 fold changes (Supplementary Table 2), gives a weighted map of the 173 genes found in BrainSpan, 141 up and 32 down (Figure 5). Cerebral organoids of this age resemble the early and mid-fetal neocortex (Camp et al. 2015; Gordon et al. 2021), and the stage test places the signature there with no information from the experiment: it is higher in the first and second trimesters (whole-brain *z* = 3.2 and 4.9), in every region in the second trimester (the cerebellum resting on one donor), and lower in the third trimester and in the child window (*z* = *−*6.4 and *−*2.4, both resting on one donor at the whole-brain level; Figure 5b). The genes ethanol raised carry this result (whole-brain *z* = 3.3 and 4.8 in the first two trimesters), while the 32 it lowered peak in no window. The score test marks 19 cells, all of them neocortical areas, in the first and second trimesters and after birth (Figure 5a): within each window, the signature stands out in the cortex rather than in subcortical structures. The marker test agrees, placing raised genes such as *MEF2C*, *RORB* and *KCNV1* among the neocortex markers of all five windows.

## 4 Discussion

VEGABrain lowers the cost of asking a spatial question of the developing human transcriptome from a small project to a few seconds in a browser. The composite enrichment layer extends the same reasoning to the unit that modern analyses actually produce—a gene set—and the on-demand mode means a user’s own list is a first-class input rather than something the tool’s authors had to anticipate. The significance tests take the last step: they turn a map into a result, by marking where and when a list stands out against random genes of the same expression level, and by naming the genes behind each area hit.

### 4.1 Relation to existing tools

Table 3 positions VEGABrain against the tools a researcher would otherwise reach for. The comparison is deliberately along axes of *what the user must supply*, not of raw capability: each of these tools is stronger than VEGABrain in its own domain, and the point is that none of them closes the particular path VEGABrain is built for—from a gene set to a scored, parcellated map of the developing brain, in a browser.

**Table 3.**
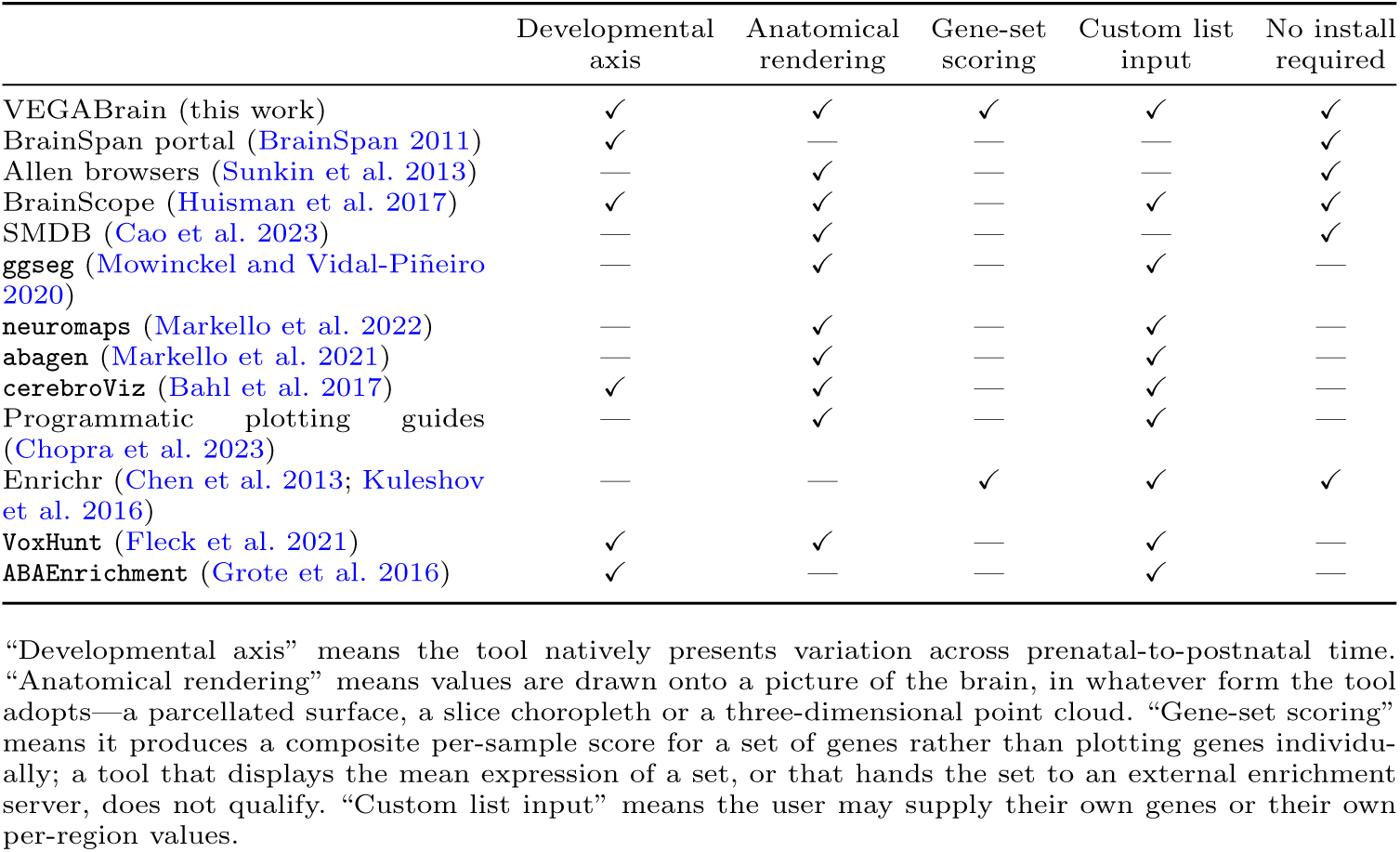
VEGABrain relative to comparable tools. ✓ = supported, — = not supported or out of scope.

|  | Developmental axis | Anatomical rendering | Gene-set scoring | Custom list input | No install required |
|---|---|---|---|---|---|
| VEGABrain (this work) | ✓ | ✓ | ✓ | ✓ | ✓ |
| BrainSpan portal (BrainSpan 2011) | ✓ | — | — | — | ✓ |
| Allen browsers (Sunkin et al. 2013) | — | ✓ | — | — | ✓ |
| BrainScope (Huisman et al. 2017) | ✓ | ✓ | — | ✓ | ✓ |
| SMDB (Cao et al. 2023) | — | ✓ | — | — | ✓ |
| ggseg (Mowinckel and Vidal-Piñeiro 2020) | — | ✓ | — | ✓ | — |
| neuromaps (Markello et al. 2022) | — | ✓ | — | ✓ | — |
| abagen (Markello et al. 2021) | — | ✓ | — | ✓ | — |
| cerebroViz (Bahl et al. 2017) | ✓ | ✓ | — | ✓ | — |
| Programmatic plotting guides (Chopra et al. 2023) | — | ✓ | — | ✓ | — |
| Enrichr (Chen et al. 2013; Kuleshov et al. 2016) | — | — | ✓ | ✓ | ✓ |
| VoxHunt (Fleck et al. 2021) | ✓ | ✓ | — | ✓ | — |
| ABAEnrichment (Grote et al. 2016) | ✓ | — | — | ✓ | — |

“Developmental axis” means the tool natively presents variation across prenatal-to-postnatal time. “Anatomical rendering” means values are drawn onto a picture of the brain, in whatever form the tool adopts—a parcellated surface, a slice choropleth or a three-dimensional point cloud. “Gene-set scoring” means it produces a composite per-sample score for a set of genes rather than plotting genes individu-ally; a tool that displays the mean expression of a set, or that hands the set to an external enrichment server, does not qualify. “Custom list input” means the user may supply their own genes or their own per-region values.

As noted in the Introduction, the portals are gene-at-a-time and the programmatic tools take a value per region as *input*, leaving the user to obtain, preprocess and map the developmental expression data themselves. cerebroViz (Bahl et al. 2017) is the closest of them in intent, having been written for spatiotemporal brain data specifically and shipping region vocabularies for BrainSpan, GTEx (GTEx Consortium 2013) and Roadmap Epigenomics (Roadmap Epigenomics Consortium et al. 2015); it nonetheless remains a drawing layer over a matrix the user provides, carries no expression data of its own, and computes no enrichment. The same qualification applies to the code-first route as a whole. Chopra et al. (2023) argue for it persuasively and lower its cost with a survey of the available packages and a generator for template plotting code, but a guide to writing the code does not remove the prerequisites the code still has: a working R or Python environment, a local copy of the expression matrix, and a mapping from tissue nomenclature onto a parcellation.

Among hosted resources, BrainScope (Huisman et al. 2017) is the nearest neighbour to VEGABrain, and the only other entry in Table 3 that is at once web-based, developmental and willing to take a user’s own gene set. The abstraction differs, and with it the question answered. BrainScope arranges the Allen adult and developing atlases as linked t-SNE maps of genes and samples, localising samples anatomically through choropleths over selected slices, and summarises a submitted set by its mean expression while delegating enrichment to Enrichr (Chen et al. 2013) and ToppGene (Chen et al. 2009). VEGABrain asks the narrower question of where a set sits in a named parcellation, window by window, and answers it with a rank-based composite score rather than a mean, drawn on the Desikan–Killiany and aseg atlases that the imaging literature already uses. SMDB (Cao et al. 2023) is likewise hosted and likewise anatomical, but for a different data type and species: it browses spatial transcriptomics sections against Allen’s mouse brain coordinate framework and reconstructs morphology in three dimensions, with no human developmental axis and no gene-set layer.

Enrichr (Chen et al. 2013; Kuleshov et al. 2016) performs the gene-set reasoning but returns ranked term tables, with no anatomical or developmental dimension. VoxHunt (Fleck et al. 2021) maps expression signatures onto developmental reference atlases, primarily to assign identity to *in vitro* organoid samples, and is an R package rather than a hosted service. VEGABrain occupies the intersection the others leave open: hosted, developmental, parcellated, and gene-set *scored*.

Testing a gene set for regional or developmental enrichment has been done before, though not in a hosted, anatomical tool. ABAEnrichment (Grote et al. 2016) tests gene sets for expression enrichment across the regions and developmental stages of the Allen atlases, BrainSpan included, as an R package that returns tables. The specificity index and the tissue-specific expression analyses built on it (Dougherty et al. 2010; Xu et al. 2014) ask, as VEGABrain’s marker test does, whether a list is over-represented among genes specific to a region and developmental period. VEGABrain combines a test of that kind with a competitive test on the composite score and with a test of developmental timing, and draws all three on the same parcellated map for any list a user pastes.

### 4.2 Limitations

Some limitations bound the interpretation of any map produced here, and the interface is designed to keep them visible rather than to smooth them over. First, donor support is small and uneven: each developmental window rests on 5 to 14 donors, and not every structure is sampled in every window. Donor counts are printed in every facet label and unsampled regions are drawn in grey rather than interpolated, and the tests of Section 2.5 leave cells with fewer than three donors untested rather than test them. Even so, the third trimester rests mostly on two donors, of 24 and 37 post-conception weeks, so a third-trimester result often reads as “near term”, and a contrast resting on few donors deserves less weight than its *q*-value alone suggests. The many-to-one assignment of BrainSpan structures to atlas parcels (Section 2.3) likewise means that neighbouring parcels sharing a structure cannot differ.

Second, the data are bulk tissue. BrainSpan samples are dissected tissue, so a regional value reflects both cell-intrinsic expression and cell-type composition: a rise in an oligodendrocyte programme across development is as consistent with more oligodendrocytes as with more transcription per cell. Single-cell atlases resolve this and are the natural next data source.

Third, ssGSEA scores are built from within-sample ranks. Equation 1 yields a value that is comparable across samples for a fixed gene set, but not across gene sets of different sizes and not on an absolute scale. Colour scales for ontology maps are therefore free rather than fixed, and a score should be read as a spatiotemporal pattern, not as a magnitude.

Fourth, the score and stage tests are competitive: they compare a list with random genes of matched expression, which ignores the correlation among the list’s own genes (Goeman and Bühlmann 2007; Wu and Smyth 2012). A coherent programme therefore passes more easily than random genes do, and because development dominates brain expression, about four in ten curated gene sets peak in some window. Each cell is reported with its effect size (*z*), which should be read alongside the mark. The marker test inherits the bulk-tissue caveat in another form: cortical areas converge after birth, so fewer area-specific genes are found and fewer postnatal areas can be tested.

Future work follows directly from these bounds: incorporating single-cell and single-nucleus developmental atlases, which would let the marker test speak of cell types rather than tissue, testing against correlation-aware nulls, and allowing user-supplied parcellations.

## 5 Conclusion

VEGABrain makes the developmental human brain transcriptome, and the activity of gene sets within it, visible anatomically without installation, coding or data preparation, and it tests submitted gene lists for where and when they are enriched against expression-matched random genes. Maps and their underlying tables are exportable, the structure-to-atlas mapping is published rather than implied, and the source is open. The tool is available at https://mcblab.github.io/VEGABrain/.

## Supplementary information

**Supplementary File 1** is the assignment of BrainSpan dissected structures to atlas parcels used by VEGABrain (Section 2.3), as a CSV table (S1 region mapping.csv) with one row per parcel–structure pair. Column region is the Desikan–Killiany or aseg parcel as named by ggseg; structure name is the BrainSpan structure whose samples are averaged into that parcel; macro regions is the macro-scale grouping (frontal, parietal, temporal or occipital lobe, subcortical, or cerebellum). A parcel fed by two structures appears on two rows (thalamus, cerebellum), and the thalamus is listed under both Thalamus and Thalamus Proper, the names it carries in different FreeSurfer aseg versions, so the table’s 29 region names denote 28 parcels.

**Supplementary Table 2** (S2 signatures.xlsx) gives every gene signature used in this work, one tab per signature: the lists of Figures 3, 4 and 5 (GOBP ASTROCYTE DEVELOPMENT, HP AUTISM and the ethanol signature of Arzua et al. 2020) and the positive controls of Section 2.5 (eight striatal markers, GOBP MYELIN ASSEMBLY, HALLMARK G2M CHECKPOINT and HALLMARK E2F TARGETS). Column gene is what was submitted to the gene-list mode, and found in BrainSpan marks the symbols present in the expression matrix, which are the only ones that enter a map or a test. The tab for Arzua et al. (2020) keeps the source table’s fold change, direction, *P* and false discovery rate, and the signed log2 fold change submitted with each gene. A README tab gives each signature’s source and the counts quoted in the text.

## Acknowledgements

We thank the Allen Institute for Brain Science and the BrainSpan consortium for making the developmental transcriptome openly available, and the maintainers of ggseg, GSVA and msigdbr, on which this work depends. The authors would like to thank NPAD / UFRN for computational resources.

## Ethics approval and consent to participate

Not applicable. This study analyses publicly released, de-identified data from the BrainSpan Developmental Transcriptome and involved no new human subjects.

## Data availability

The expression data analysed here are the BrainSpan Developmental Transcriptome RNA-Seq Gencode v10 summarised-to-genes release (BrainSpan 2011; Li et al. 2018), publicly available at https://www.brainspan.org. Gene sets were obtained from MSigDB through the msigdbr R package. The derived per-region, per-window score tables served by VEGABrain are downloadable as CSV directly from the application.

## Code availability

VEGABrain is open source at https://github.com/MCBLab/VEGABrain. The application is deployed at https://mcblab.github.io/VEGABrain/; its public HTTP API is documented in the repository.

## Author contributions

IM: Formal analysis, Methodology, Visualization, Writing – review and editing. JVFC: Formal analysis, Writing – review and editing. DMC: Conceptualization, Data curation, Investigation, Supervision, Writing – original draft, Writing – review and editing.

## References

Arzua T, Yan Y, Jiang C, et al (2020) Modeling alcohol-induced neurotoxicity using human induced pluripotent stem cell-derived three-dimensional cerebral organoids. Translational Psychiatry 10(1):347. 10.1038/s41398-020-01029-4

Ashburner M, Ball CA, Blake JA, et al (2000) Gene Ontology: tool for the unification of biology. Nature Genetics 25(1):25–29. 10.1038/75556

Bahl E, Koomar T, Michaelson JJ (2017) cerebroViz: an R package for anatomical visualization of spatiotemporal brain data. Bioinformatics 33(5):762–763. 10.1093/bioinformatics/btw726

Barbie DA, Tamayo P, Boehm JS, et al (2009) Systematic RNA interference reveals that oncogenic KRAS-driven cancers require TBK1. Nature 462(7269):108–112. 10.1038/nature08460

Benjamini Y, Hochberg Y (1995) Controlling the false discovery rate: a practical and powerful approach to multiple testing. Journal of the Royal Statistical Society: Series B (Methodological) 57(1):289–300. 10.1111/j.2517-6161.1995.tb02031.x

BrainSpan (2011) BrainSpan: Atlas of the Developing Human Brain. https://www.brainspan.org, RNA-Seq Gencode v10 summarized to genes; funded by ARRA awards 1RC2MH089921-01, 1RC2MH090047-01 and 1RC2MH089929-01. Accessed 2026-08-18

Camp JG, Badsha F, Florio M, et al (2015) Human cerebral organoids recapitulate gene expression programs of fetal neocortex development. Proceedings of the National Academy of Sciences 112(51):15672–15677. 10.1073/pnas. 1520760112

Cao R, Ling Y, Meng J, et al (2023) SMDB: a spatial multimodal data browser. Nucleic Acids Research 51(W1):W553–W559. 10.1093/nar/gkad413

Chen EY, Tan CM, Kou Y, et al (2013) Enrichr: interactive and collaborative HTML5 gene list enrichment analysis tool. BMC Bioinformatics 14(1):128. 10.1186/1471-2105-14-128

Chen J, Bardes EE, Aronow BJ, et al (2009) ToppGene Suite for gene list enrichment analysis and candidate gene prioritization. Nucleic Acids Research 37(Web Server issue):W305–W311. 10.1093/nar/gkp427

Chopra S, Labache L, Dhamala E, et al (2023) A practical guide for generating reproducible and programmatic neuroimaging visualizations. Aperture Neuro 3. 10.52294/001c.85104

Desikan RS, Ségonne F, Fischl B, et al (2006) An automated labeling system for subdividing the human cerebral cortex on MRI scans into gyral based regions of interest. NeuroImage 31(3):968–980. 10.1016/j.neuroimage.2006.01.021

Dolgalev I (2026) msigdbr: MSigDB gene sets for multiple organisms in a tidy data format. https://igordot.github.io/msigdbr/, R package version 26.1.0

Dougherty JD, Schmidt EF, Nakajima M, et al (2010) Analytical approaches to RNA profiling data for the identification of genes enriched in specific cells. Nucleic Acids Research 38(13):4218–4230. 10.1093/nar/gkq130

Fischl B, Salat DH, Busa E, et al (2002) Whole brain segmentation: automated labeling of neuroanatomical structures in the human brain. Neuron 33(3):341–355. 10.1016/s0896-6273(02)00569-x

Fleck JS, Sanchís-Calleja F, He Z, et al (2021) Resolving organoid brain region identities by mapping single-cell genomic data to reference atlases. Cell Stem Cell 28(6):1148–1159.e8. 10.1016/j.stem.2021.02.015

Gillespie M, Jassal B, Stephan R, et al (2022) The Reactome pathway knowledge-base 2022. Nucleic Acids Research 50(D1):D687–D692. 10.1093/nar/gkab1028

Goeman JJ, Bühlmann P (2007) Analyzing gene expression data in terms of gene sets: methodological issues. Bioinformatics 23(8):980–987. 10.1093/bioinformatics/btm051

Gordon A, Yoon SJ, Tran SS, et al (2021) Long-term maturation of human cortical organoids matches key early postnatal transitions. Nature Neuroscience 24(3):331–342. 10.1038/s41593-021-00802-y

Grote S, Prüfer K, Kelso J, et al (2016) ABAEnrichment: an R package to test for gene set expression enrichment in the adult and developing human brain. Bioinformatics 32(20):3201–3203. 10.1093/bioinformatics/btw392

GTEx Consortium (2013) The Genotype-Tissue Expression (GTEx) project. Nature Genetics 45(6):580–585. 10.1038/ng.2653

Hawrylycz MJ, Lein ES, Guillozet-Bongaarts AL, et al (2012) An anatomically comprehensive atlas of the adult human brain transcriptome. Nature 489(7416):391–399. 10.1038/nature11405

Huisman SMH, van Lew B, Mahfouz A, et al (2017) BrainScope: interactive visual exploration of the spatial and temporal human brain transcriptome. Nucleic Acids Research 45(10):e83. 10.1093/nar/gkx046

Hänzelmann S, Castelo R, Guinney J (2013) GSVA: gene set variation analysis for microarray and RNA-seq data. BMC Bioinformatics 14(1):7. 10.1186/1471-2105-14-7

Kanehisa M, Goto S (2000) KEGG: Kyoto Encyclopedia of Genes and Genomes. Nucleic Acids Research 28(1):27–30. 10.1093/nar/28.1.27

Kang HJ, Kawasawa YI, Cheng F, et al (2011) Spatio-temporal transcriptome of the human brain. Nature 478(7370):483–489. 10.1038/nature10523

Kuleshov MV, Jones MR, Rouillard AD, et al (2016) Enrichr: a comprehensive gene set enrichment analysis web server 2016 update. Nucleic Acids Research 44(W1):W90–W97. 10.1093/nar/gkw377

Köhler S, Gargano M, Matentzoglu N, et al (2021) The Human Phenotype Ontology in 2021. Nucleic Acids Research 49(D1):D1207–D1217. 10.1093/nar/gkaa1043

Lau C, Ng L, Thompson C, et al (2008) Exploration and visualization of gene expression with neuroanatomy in the adult mouse brain. BMC Bioinformatics 9(1):153. 10.1186/1471-2105-9-153

Li M, Santpere G, Imamura Kawasawa Y, et al (2018) Integrative functional genomic analysis of human brain development and neuropsychiatric risks. Science 362(6420):eaat7615. 10.1126/science.aat7615

Liberzon A, Subramanian A, Pinchback R, et al (2011) Molecular Signatures Database (MSigDB) 3.0. Bioinformatics 27(12):1739–1740. 10.1093/bioinformatics/btr260

Liberzon A, Birger C, Thorvaldsdóttir H, et al (2015) The Molecular Signatures Database hallmark gene set collection. Cell Systems 1(6):417–425. 10.1016/j.cels.2015.12.004

Markello RD, Arnatkeviciute A, Poline JB, et al (2021) Standardizing workflows in imaging transcriptomics with the abagen toolbox. eLife 10:e72129. 10.7554/elife.72129

Markello RD, Hansen JY, Liu ZQ, et al (2022) neuromaps: structural and functional interpretation of brain maps. Nature Methods 19(11):1472–1479. 10.1038/s41592-022-01625-w

Miller JA, Ding SL, Sunkin SM, et al (2014) Transcriptional landscape of the prenatal human brain. Nature 508(7495):199–206. 10.1038/nature13185

Mowinckel AM, Vidal-Piñeiro D (2020) Visualization of brain statistics with R packages ggseg and ggseg3d. Advances in Methods and Practices in Psychological Science 3(4):466–483. 10.1177/2515245920928009

Parikshak NN, Luo R, Zhang A, et al (2013) Integrative functional genomic analyses implicate specific molecular pathways and circuits in autism. Cell 155(5):1008–1021. 10.1016/j.cell.2013.10.031

R Core Team (2026) R: A language and environment for statistical computing. https://www.R-project.org/, Version 4.6.0

Ritchie ME, Phipson B, Wu D, et al (2015) limma powers differential expression analyses for RNA-sequencing and microarray studies. Nucleic Acids Research 43(7):e47. 10.1093/nar/gkv007

Roadmap Epigenomics Consortium, Kundaje A, Meuleman W, et al (2015) Integrative analysis of 111 reference human epigenomes. Nature 518(7539):317–330. 10.1038/nature14248

Rowitch DH, Kriegstein AR (2010) Developmental genetics of vertebrate glial-cell specification. Nature 468(7321):214–222. 10.1038/nature09611

Schloerke B, Allen J (2026) plumber: An API generator for R. https://www.rplumber.io, R package version 1.3.3

Subramanian A, Tamayo P, Mootha VK, et al (2005) Gene set enrichment analysis: A knowledge-based approach for interpreting genome-wide expression profiles. Proceedings of the National Academy of Sciences 102(43):15545–15550. 10.1073/pnas.0506580102

Sunkin SM, Ng L, Lau C, et al (2013) Allen Brain Atlas: an integrated spatio-temporal portal for exploring the central nervous system. Nucleic Acids Research 41(D1):D996–D1008. 10.1093/nar/gks1042

The Gene Ontology Consortium, Aleksander SA, Balhoff J, et al (2023) The Gene Ontology knowledgebase in 2023. Genetics 224(1):iyad031. 10.1093/genetics/iyad031

Wickham H (2016) ggplot2: Elegant Graphics for Data Analysis. Use R!, Springer-Verlag New York, 10.1007/978-3-319-24277-4

Wilkinson MD, Dumontier M, Aalbersberg IJ, et al (2016) The FAIR guiding principles for scientific data management and stewardship. Scientific Data 3(1):160018. 10.1038/sdata.2016.18

Willsey AJ, Sanders SJ, Li M, et al (2013) Coexpression networks implicate human midfetal deep cortical projection neurons in the pathogenesis of autism. Cell 155(5):997–1007. 10.1016/j.cell.2013.10.020

Wu D, Smyth GK (2012) Camera: a competitive gene set test accounting for inter-gene correlation. Nucleic Acids Research 40(17):e133. 10.1093/nar/gks461

Xu X, Wells AB, O’Brien DR, et al (2014) Cell type-specific expression analysis to identify putative cellular mechanisms for neurogenetic disorders. The Journal of Neuroscience 34(4):1420–1431. 10.1523/JNEUROSCI.4488-13.2014

Yi M, Nissley DV, McCormick F, et al (2020) ssGSEA score-based Ras dependency indexes derived from gene expression data reveal potential Ras addiction mechanisms with possible clinical implications. Scientific Reports 10(1):10258. 10.1038/s41598-020-66986-8

Zhu Y, Sousa AMM, Gao T, et al (2018) Spatiotemporal transcriptomic divergence across human and macaque brain development. Science 362(6420):eaat8077. 10.1126/science.aat8077

